# Sound-Enhanced Sleep Depth Reduces Traumatic Brain Injury Damage and Sequelae and Supports Microglial Response

**DOI:** 10.1101/2025.01.21.634054

**Authors:** Carlos G. Moreira, Adrian Müllner, Meltem Gönel, Pascal Hofmann, Filipe Teixeira, Rosa C. Paolicelli, Inês Dias, Sergio I. Nemirovsky, Christian R. Baumann, Daniela Noain

## Abstract

Traumatic brain injury significantly reduces the quality of life for millions of survivors worldwide, with no established treatments currently available. High slow-wave activity (SWA) sleep immediately after rodent TBI improves posttraumatic outcomes. However, pharmacological SWA-enhancing strategies are hindered by severe specificity and scalability issues that prevent it from effectively reaching clinical implementation. Alternatively, closed-loop auditory stimulation (CLAS) of sleep slow waves offers specific SWA enhancement with high translational potential. Our present results demonstrate that up-phase-targeted CLAS (upCLAS)-mediated SWA enhancement reduced diffuse axonal injury, decreased demyelination, and preserved cognitive ability in TBI rats. The alleviated posttraumatic phenotype was associated with increased microglia response, likely mediating CLAS’ neuroprotective effect in the acute injury phase. Auditory-enhanced SWA may thus constitute a novel noninvasive neuroprotective therapy preventing TBI sequelae via boosted cellular response to tissue damage.

## Introduction

Every year, 69 million people worldwide are estimated to sustain a traumatic brain injury (TBI), and nearly half of these individuals report at least three persistent posttraumatic symptoms, including cognitive deficits^1–3^. One common TBI-associated neuropathology is diffuse axonal injury (DAI). Damage incurred by brain white matter from trauma forces causes complex cytoskeletal changes that lead to rapid accumulation of proteinaceous products, such as amyloid precursor protein (APP), in axonal swellings or bulbs^4^. In severe cases, DAI can result in demyelination, ultimately ending in axotomy and cell death^5^, leading to significant functional impairments in TBI patients. As a first response to mitigate brain damage after TBI, acute recruitment of microglia – the central nervous system resident innate immune cells – plays a critical role, detecting and rapidly responding to tissue injury and often acting to remove cellular debris^6,7^. In fact, defective microglia response in TBI may be predictive of cognitive deficits^8^.

It is known that deep sleep supports the accumulation and removal of toxic proteinaceous metabolites from the brain via regulation of release and clearance processes, which rather straightforwardly explains the impact of sleep modulation onto soluble protein content^9–12^. However, whether such sleep-regulated pathways mediate the accumulation/removal of larger protein complexes and/or cellular debris after eg. TBI is less certain, suggesting that other processes downstream of sleep may be relevant. Glia neuroinflammatory response, on the other hand, has been shown to play a positive role in the acute phase after trauma^13,14^, indicating that microglia could initially mediate neuroprotective processes after TBI. Strikingly, direct links between glial cells and processes driving the sleep-wake cycle have been identified recently^15–17^, enabling the notion that modulation of sleep/circadian cycle could alter microglial responses in the context of disease.

Thus, by interacting with potentially neuroprotective brain processes triggered upon tissue damage, sleep modulation has been conceptualized as a potential therapy after TBI^18–20^. We have shown that pharmacologically increased slow wave activity (SWA) during the first days after rodent TBI significantly reduces APP accumulation and preserves posttraumatic cognitive ability^21^. However, pharmacological approaches to enhance SWA alter both the cyclic transitions between sleep stages and physiological non-rapid eye movement (NREM) sleep^22,23^, thus do not constitute a sleep architecture-preserving methodology. In addition, they lack the specificity and innocuousness required for scaled clinical implementation. Instead, an ideal therapeutic strategy for TBI victims should preserve sleep architecture and enhance SWA via a nonpharmacological, noninvasive method, such as up-phase-targeted closed-loop auditory stimulation (upCLAS)^24^.

In this study, we delivered upCLAS to rats immediately after TBI to: (i) determine the efficacy, safety, specificity, and stability of upCLAS as SWA modulator after TBI; (ii) demonstrate the critical role of increased SWA in reducing TBI-associated neuropathology and posttraumatic cognitive deficits; and iii) explore the extent of a potential sleep-mediated cellular immune response in the brain after TBI.

## Materials and methods

### Animals, surgeries, and husbandry

We used 20 young-adult male Sprague-Dawley rats (Charles River, Italy) weighing 250– 300g, and group-housed them in standard individually ventilated cages (T2000) prior to interventions. We surgically implanted electrodes in all animals for continuous recording of electroencephalography and electromyography (EEG/EMG) as described previously^25^. Briefly, we inserted four stainless steel miniature screws (Hasler, Switzerland), one pair for each hemisphere, bilaterally into the rats’ skulls following specific stereotaxic coordinates: the anterior electrodes were implanted 3mm posterior to bregma and 2mm lateral to the midline, and the posterior electrodes 6mm posterior to bregma and 2mm lateral to the midline. For monitoring muscle tone, we inserted a pair of gold wires as EMG electrodes into the rats’ neck muscles. All electrodes were connected to stainless steel wires, further connected to a headpiece (Farnell, #M80-8540842, Switzerland), and fixed to the skull with dental cement. We performed all surgical procedures under deep anaesthesia by inhalation of isoflurane (4.5% for induction, 2.5% for maintenance) and subsequent analgesia with buprenorphine (s.c. 0.05 mg/kg). Following surgery, we housed the animals in pairs for a minimum of 14 days for recovery, with food and water available ad libitum, and handled them daily for postoperative monitoring, body-weight check, and familiarization with the experimenter. The animal-room temperature was maintained at 22–23 °C, and animals were kept on a 12h light–dark cycle. All procedures were approved by the Veterinary Office of the Canton Zurich (license ZH231/2015) and conducted in accordance with national and institutional regulations for care and use of laboratory animals.

### Experimental design

We performed the experiment in multiple batches, allocated evenly across 3 experimental groups: non-TBI rats (n = 8) underwent no traumatic brain injury (TBI) and posttraumatic mock closed-loop auditory stimulation (CLAS) of slow waves (events flagged but no sound delivered); TBI mockCLAS rats (n = 7) underwent TBI and posttraumatic mock CLAS of slow waves; and TBI upCLAS rats (n = 8) received both TBI and up-phase-targeted CLAS of slow waves (pink noise triggers delivered to the slow waves’ up-phase) (**Figure 1a**). Following the first novel-object recognition test (NORT), we transferred the rats to individual custom-made acrylic-glass cages (26.5 × 42.5 × 43.5 cm). Each cage was positioned inside a sound-attenuated chamber, built in-house, for 24-h undisturbed EEG/EMG baseline (BL) recording^24^. The day after, rats underwent closed-skull TBI induction or sham surgery. On the following day, we initiated a 5-day auditory stimulation protocol^24^. CLAS was stopped at the end of the 5^th^ day, and the animals were recorded for 2 more days without stimulation, to assess potential carry-over effects. Duration of treatment was determined by our previous studies^25^, in which we detected a cognitive deficit at 7 days post-TBI, leaving a possible window for interventions of 5 days. Two weeks following sham or TBI induction, the rats were again tested in the NORT, and 28 days after sham or TBI induction, we euthanized the rats and harvested their brains for histopathological determinations. A minimum of 4 and a maximum of 7 animals per group were analysed for each measure, as reported in detail in the figure legends. Exclusions of data in the different outcomes were related to low EEG/EMG signal quality or loss (system crashes), non-engaging animal in the behavioural task (median split discussed for NORT, below), and lack of tissue integrity or availability (APP, MBP, IBA1).

**Figure 1.**
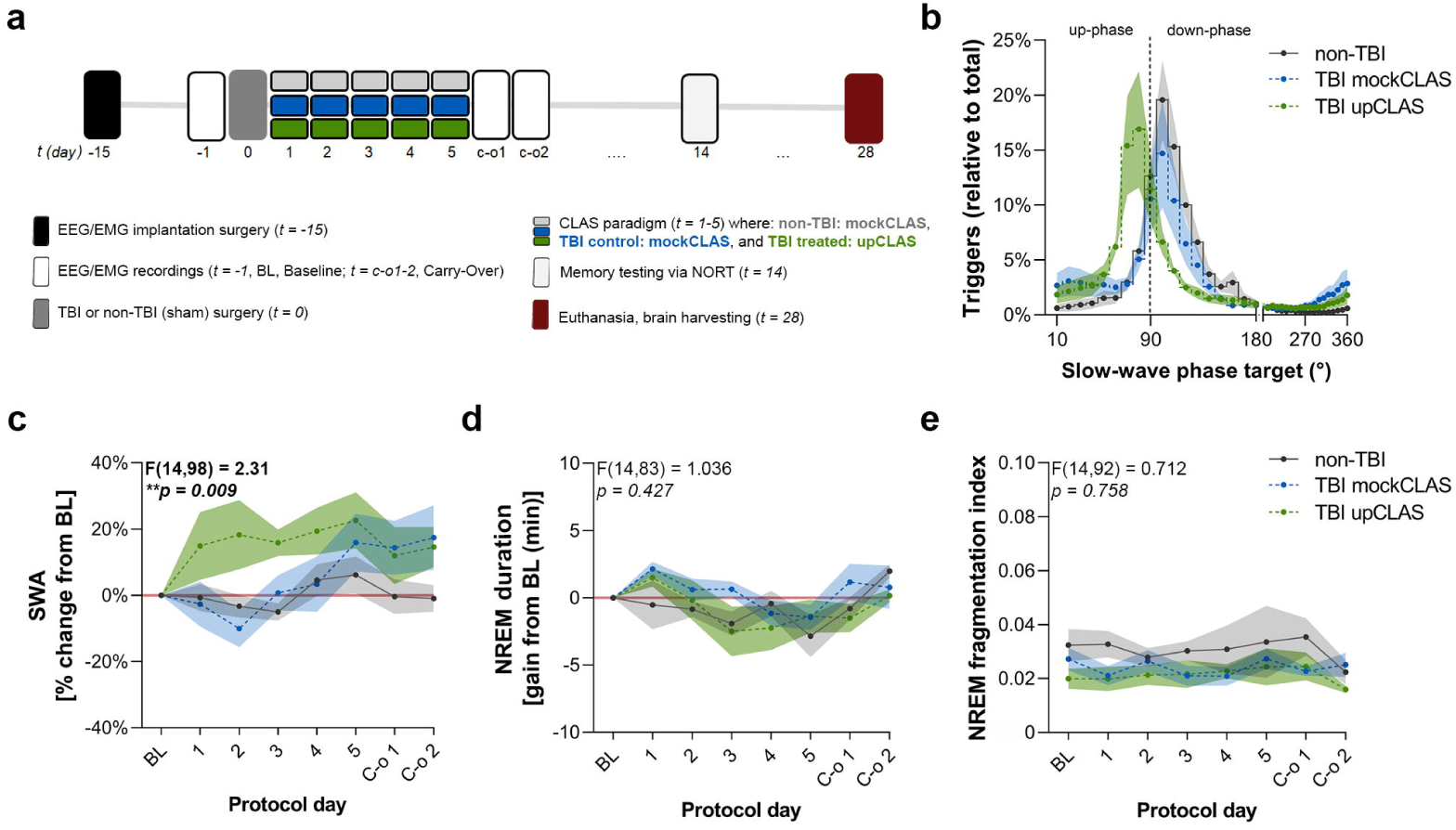
upCLAS boosts SWA without influencing NREM sleep architecture. **a)** Experimental design depicting the succession of events per protocol day (*t*) for all animals in each group. Non-TBI (grey) and TBI mockCLAS ( blue) groups received no sound stimulation during the CLAS paradigm, but the corresponding stimulation events – targeted to the down-phase – were flagged; whereas the TBI upCLAS (green) group received sound triggers targeted to the up-phase of slow waves during NREM sleep. **b)** Percentage of target phase distribution relative to total targets for each group, demonstrating target accuracy for the up-phase of slow waves in the TBI upCLAS group. **c)** SWA (two-way ANOVA, treatment × TBI interaction; F(14, 98)□=□2.31, \*\**p* = 0.009), **d)** NREM sleep duration (two-way ANOVA, treatment × TBI interaction; F(14, 83)□=□1.036, *p* = 0.427) and **e)** NREM sleep fragmentation index (two-way ANOVA, treatment × TBI interaction; F(14, 92)□=□0.712, *p* = 0.758) at baseline (BL), during the 5□days of modulation (1–5), and the two subsequent observation days assessing carry-over effects (C-o 1, C-o 2) in non-TBI (n = 6), TBI mockCLAS (n = 7), and TBI upCLAS (n = 6) rats. SWA: slow-wave activity; NREM: non-rapid eye movement sleep; upCLAS: up-phase targeted closed-loop auditory stimulation; mockCLAS: flagging of triggers’ targets without sound delivery; TBI: traumatic brain injury; non-TBI: sham operated rats.

### Novel object recognition test

To examine declarative memory performance, we used the NORT as described previously^21,26^. Briefly, the test consisted of two phases: habituation and testing. In the habituation phase, the animal was presented for 10=min of exploration with two equal objects; these became the familiar objects. Three hours later, the testing phase consisted of reintroducing the animal to the context and presenting both an identical copy of the familiar object, to avoid olfactory cues, and a novel object. We left the animal free to explore both objects for 5=minutes. The testing phase was acquired and analysed using dedicated software (Ethovision, Noldus, Germany). We extracted the time spent exploring the two objects, defined as biting or sniffing at distance of <2=cm, and calculated the recognition index, defined as time spent exploring the novel object relative to the total exploration time.

### Traumatic brain injury induction

TBI induction was conducted as described previously^21,25,27^. Briefly, rats were deeply anesthetized under 2.5% isoflurane and positioned on a foam platform (L□×□W□×□T: 17□×□9□×□10□cm, stiffness: 2.84□Newton/cm). Their heads were closely bordered bilaterally with two smooth wooden walls for correct positioning during trauma induction. A 0.5–0.7□cm scalp incision over the midline in the frontal head area was made just anterior to the implanted EEG/EMG headset. After exposing the skull, the impact area (2□mm anterior to bregma, over the midline) was marked and a 1 mm-thick metal plate (1□cm circumradius) was placed over the exposed target area to prevent bone fractures. A 2500□g stainless steel rod of a total length of 100□cm, diameter 2□cm, with a flat pointed silicon tip (diameter 1□mm, total tip length 4□cm) was used as falling weight. It was mounted in a slide and held by a stand at an angle of 70 degrees. The metal rod was positioned precisely over the selected point of injury, elevated to a height of 25□cm, and released by pushing a button deactivating a magnet that held the metal rod in place. Non-TBI animals underwent the exact same procedures except for the injury. Subsequently, the skin was closed and disinfected. Animals returned to their home cages and were monitored continuously for at least 1□h until they had fully recovered from sedation and normal home cage behaviour was observed.

### Full body perfusion and brain sectioning

Twenty-eight days after sham or TBI procedures, we euthanized all rats by transcardiac perfusion^21,25^. Briefly, we exsanguinated the animals using ice-cold phosphate-buffered saline (PBS) followed by perfusion of freshly prepared ice-cold 4% paraformaldehyde (PFA) (MilliporeSigma, Germany). Immediately after fixation, we carefully harvested the brains, postfixed them for 12 hours in 4% PFA + 15% sucrose and further dehydrated them for 48– 72 hours in 30% sucrose. We then quick-froze the brains in dry ice and stored them at -80 °C until further use. Later, full-brain serial sectioning was performed in stereological fashion at 40 μm thickness using a freezing-stage-equipped microtome (Leica).

### EEG/EMG recording and pre-processing

To verify the effect of CLAS on EEG spectra, we conducted bilateral tethered EEG/EMG recordings in differential mode for 24 h to serve as BL and throughout all the subsequent 7 days of the protocol (5 days of stimulation, days 1–5 + 2 carry-over (c-o) days, days c-o 1 and c-o 2), applying our runtime stimulation paradigm in 5 or 6 freely moving animals simultaneously. We acquired data using a multichannel neurophysiology recording system (Tucker Davis Technologies, TDT, USA). We sampled all EEG/EMG signals at 610.35 Hz, amplified them (PZ5 NeuroDigitizer preamplifier, TDT, USA) after applying an anti-aliasing low-pass filter (45% of sampling frequency), synchronously digitized them (RZ2 BIOAMP processor, TDT, USA), recorded them using SYNAPSE software (TDT, USA), and stored them locally (WS-8 workstation, TDT, USA). We filtered real-time EEG between 0.1 and 36.0 Hz (2nd order biquad filter, TDT, USA), and EMG between 5.0 and 525.0 Hz (2nd order biquad filter and 40-dB notch filter centred at 50 Hz, TDT, USA), and fed the signals to real-time detection algorithms for non-rapid eye movement (NREM) sleep staging and phase detection.

### Online NREM sleep staging and phase-locked auditory closed-loop stimulation of sleep slow waves

Parallel rule-based NREM sleep (NREMS) staging and phase detection features were run continuously alongside EEG/EMG recordings so that sound triggers were presented in real time at every instance the stimulatory truth function compounding these features was reached. For online sleep staging, a nonlinear classifier compounded two major decision nodes: power in EEG and power in EMG. Briefly, we computed high-beta (20–30 Hz) and delta (0.5–4 Hz) bands’ root mean square (rms) on a sliding window of 1 s using an algorithm written in RPvdsEx (Real-time Processor visual design studio, TDT, USA). Once the rmsdelta/rmshigh beta ratio, hereinafter referred as the NREMratio, crossed a threshold individually identified during the BL recording, we further compared EMG rms to a threshold, also defined during BL, to rule out movement artefacts. The NREMratio and the EMG power thresholds for online NREM staging during auditory stimulation, suggestive of sustained SWS during NREM, were extracted individually and immediately after the BL recording: 24 h EEG/EMG data was scored automatically using SPINDLE (Sleep phase identification with neural networks for domain-invariant learning)^28^ and fed to a custom-written MATLAB (ver. R2022b) script. In short, a strict estimate of NREMratio threshold during NREM was established as +1.0 SD over the mean, representing the 84.1% percentile, of the NREMratio of all NREMS epochs during BL. Similarly, EMG power in NREMS was delimited to values - 1.0 SD below the mean of the EMG rms values (threshold at 15.9% percentile) during offline-scored NREMS. These two values marked the transition into consolidated NREMS in each subject during online NREMS staging and were introduced to a customized SYNAPSE® (TDT, USA) project. For phase-targeted auditory stimulation of slow waves, SYNAPSE® combines the online NREMS staging feature with a phase detector. Briefly, a runtime very narrow bandpass filter (TDT, USA) for EEG phase detection isolated the 1-Hz component for phase targeting of slow waves, approximately 1.35 Hz in rats^29^, in each subjects’ left EEG channel. A threshold rule was added to avoid stimulation during oscillatory activity at very low amplitude. We predetermined slow wave’s 0° as the rising zero crossing, 90° as the positive peak, and 270° as the slow wave’s trough. At every identified positive zero-crossing on the filtered signal, the phase detector reset to 0° and calculated any of the selected target phases from the number of samples elapsed since the zero crossing. This method offers the chance to recognize slow waves consistently across conditions independently of the target phase. We divided the animals into 2 phase-targeted stimulation approaches: up-phase stimulation targeting 65° and mock stimulation arbitrarily flagging slow waves at 90° with no delivery of sound. A sound trigger was sent within 8 ms (RX8 MULTI-I/O processor, TDT) upon validation of truth-value for NREMratio, EMG power and phase target criteria. Nonarousing stimuli consisted of clicks of pink 1/f noise (30 ms duration, 35 dB SPL, 2 ms rising and falling slopes) in free field conditions from built-in speakers (MF1 Multi-field magnetic speakers, TDT) on top of the stimulation chamber 50 cm above the centre of the floor area. The protocol for the mock condition was similar to up-phase stimulation, but the sound was muted. All triggers were time-flagged for offline analysis. Throughout all days of stimulation, we used a video system to sporadically control for stimuli-evoked arousals or reflexes.

### EEG/EMG scoring

We scored all recording files using the SPINDLE online computational tool for animal sleep data (https://sleeplearning.ethz.ch/)^28^. In short, European Data Format (.edf) files, consisting of 2 parietal EEG channels and 1 nuchal EMG channel, were uploaded to SPINDLE to retrieve vigilance states with 4-second epoch resolution. The algorithm classified 3 vigilance states: wakefulness, NREMS, and REMS. Additionally, unclear epochs or interfering signals were labelled as artifacts in wakefulness, NREMS, and REMS. Wakefulness was defined by high or phasic EMG activity for more than 50% of the epoch duration and low amplitude but high frequency EEG. NREMS was characterized by reduced or no EMG activity, increased EEG power in the frequency band < 4 Hz, and the presence of slow waves. REMS was defined by high theta power (6 – 9 Hz frequency band) and low muscle tone.

### Post-processing of EEG and EMG

Time spent in NREMS was determined as an absolute number of minutes for BL or stimulation period. For the same days, we extracted measures of global spectral responses in the delta frequency, by processing the left-hemisphere EEG signal with a custom MATLAB routine (ver. R2016b). Briefly, we removed artifacts by detecting clipping events (15 adjacent raw EEG samples of approximately 60 ms within 55 units of the amplifier maximum or minimum), followed by a 3-point moving average (to remove frequencies greater than 80 Hz). Subsequently, we applied a basic Fermi window function, f(n)=(1+e^((5-n⁄50)))^((-1)), to gradually attenuate the first and last 2 s of each signal recorded (n = 600), and resampled the EEG signal at 300 Hz. Next, we filtered the signal between 0.5 Hz and 48 Hz using low- and high-pass zero-phased equiripple FIR filters (Parks-McClellan algorithm; applied in both directions (filtfilt); order_high = 1880, order_low = 398; -6 dB (half-amplitude) cut-off: high pass = 0.28 Hz, low-pass = 49.12 Hz). The signal was visually inspected for any regional artifacts (2-hour sliding window) not detected during automatic scoring: within scored NREMS, brief portions (< 10 sample-points at 300 Hz) of signal >± 8 × interquartile range were reconstructed by piecewise cubic spline interpolation from neighbouring points. We performed spectral analysis of consecutive 4 s epochs (FFT routine, Hamming-window, 2 s overlap, resolution of 0.25 Hz), and normalized the power estimate of each frequency bin in relation to the total spectral power (0.5 – 30 Hz). Additionally, we calculated hourly SWA (0.5–4 Hz) as the mean spectral power of equally sized NREMS-epoch bins (12 bins during light period and 6 bins during dark period), using the digital filters mentioned above (order_high = 3758 and order_low = 3861) normalized by the hourly total power (0.5–30 Hz). Hourly values were extrapolated from this curve.

### Immunohistochemistry of markers of structural brain integrity

#### Amyloid precursor protein (APP) axonal bulbs

Diffuse axonal injury (DAI) is evidenced by APP immunoreactive axonal accumulation^30^. We selected the corpus callosum for its especially high vulnerability to shearing, straining, and compression TBI forces, similar to that of the cerebral cortex. In fact, in parasagittal white matter of the cerebral cortex and corpus callosum, large numbers of damaged axonal projections are found after TBI due to deformation and disruption of the neurofilament subunits within the intra-axonal and intracellular cytoskeleton^31,3231,3261,6231,3231,3231,32^. To evaluate the effect of CLAS on DAI in white matter tracts, we performed unbiased immunohistochemistry and stereology to quantify the numbers of APP-stained axonal swellings or bulbs on the anterior portion of the corpus callosum in 40 μm coronal brain sections. Briefly, all sections were immersed in 10 mM citrate buffer and placed in water bath at 80°C for 30 min for antigen retrieval and then allowed to cool to room temperature for 10 min. Next, sections were washed in PBS, and endogenous peroxidase activity was quenched with 0.6% hydrogen peroxide in PBS for 30=min. After blocking of nonspecific binding sites with 5% normal goat serum (Sigma-Aldrich, Germany) in tris-buffered saline (TBS) containing 0.25% Triton-X (TBST) for 60=min, the slides were incubated with a primary beta-amyloid polyclonal antibody (cat#: 512700 (CT695), Invitrogen, USA) in a 1:1000 dilution in TBST at 4°C for 48 hours. After TBS/TBST washings, the sections were incubated with goat anti-rabbit secondary antibody (biotinylated, BA-1000, VectorLaboratories, USA) in a 1:500 dilution in TBST with 2% normal goat serum at room temperature for 4 hours. We then incubated in ABC-Elite kit (PK6105, VectorLaboratories, USA) in a 1:100 dilution in TBST on ice for 1 h. Lastly, we washed all sections in tris-buffer (TB) for 10 min and developed the signal in 0.025% 3,3′-diaminobenzidine + 0.01% H_2_O_2_ in TB with shaking for 8–10 minutes. After completion, sections were washed in PBS, mounted on glass slides, and dried overnight. Once dry, the slides were dipped for 3=min each in 70%, 95%, 100%, and 100% ethanol solutions, followed by Roticlear (Carl Roth, Germany). Finally, the slides were cover-slipped with Rotimount (Carl Roth, Germany) for microscopy and stereological quantification.

#### Myelin integrity

In TBI, myelin is damaged through both direct axonal injury and indirect secondary injuries, such as DAI, astroglial damage, autoimmunity, and cell body injury^33^. To assess whether CLAS contributes to preventing further pathophysiological mechanisms other than or downstream of DAI, we assessed myelin integrity of axonal projections in the corpus callosum with specific immunohistochemistry against myelin basic protein (MBP), which constitutes one of the most abundant proteins in the myelin sheath. We used a similar DAB immunohistochemistry protocol as the one described above for APP detection, except for the antigen retrieval step. Briefly, non-TBI and TBI brains were cryosectioned into 40 µm coronal sections of the rostral portion of the corpus callosum and then DAB-immunostained against MBP using a mouse monoclonal anti-MBP primary antibody ((F-6) sc-271524, Santa Cruz Biotechnology, USA) in a 1:500 dilution in TBST followed by a biotinylated goat anti-mouse secondary antibody (A16076, Invitrogen, USA) in a 1:1000 dilution in TBST. The stained tissue was cover-slipped for microscopy and optical density quantification.

### Quantification of DAI and demyelination measures

#### Stereological estimates of APP immunoreactive axonal bulbs in the corpus callosum

All stereological estimations were performed by an experimenter blinded to the experimental group. The optical fractionator technique was used to count a systematic random sample of positively stained axonal bulbs within the anterior part of the corpus callosum in collections from two defined adjacent brain regions containing 6 sections each. In brief, we outlined the corpus callosum from each coronal section at low power (4×) using an Axio Imager M2 microscope (Karl Zeiss, Jena, Germany), fitted with a mechanical specimen stage (75x50 mot, CAN (D) for Axio Imager). Next, we systematically counted bulbs at high magnification (63×, oil immersion), of sites within the region examined (Stereo Investigator 10.5 software, MBF Bioscience, USA). To quantify APP, a 200 × 200 μm sampling grid and a 100 × 100 μm counting frame were used. Nonaxonal DAB-stained structures (e.g., DAB-positive red blood cells) were not selected. These parameters allowed reasonable accuracies in cell counts, as determined by Gundersen’s coefficients of error between 5% and 15%.

#### Optical density of MBP staining in the corpus callosum

We used the bright field microscope Axio Imager M2 (Karl Zeiss, Jena, Germany) to acquire images from the corpus callosum in both hemispheres. The region of interest was the corpus callosum in the hemisphere with the highest signal intensity for quantification. We obtained full-section images with a 2.5 × objective. All images were processed in ImageJ (Fiji) using the Hematoxilin-Eosin Diaminobencidine (H&E DAB) vector. We then measured the average colour intensity, which is proportional to the concentration of the staining in the corpus callosum. To correct for background noise, we subtracted the intensity in the striatum. Following the Lambert-Beer law, we calculated the optical density with the formula

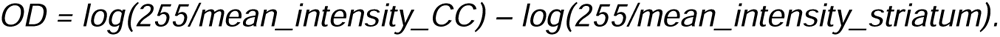

### Microglia profiling

Brain slices were permeabilized with 0.5% Triton X-100 (Sigma-Merck, #X100) in PBS for 1.5h at RT and blocking was performed with 2% BSA (VWR, #9048-46-8) in permeabilization buffer (1h at RT). Primary antibody against Ionized calcium-binding adaptor molecule 1 (IBA1, 1:1000, FUJIFILM Wako, #019-19741) was diluted in blocking buffer and slices were incubated overnight at 4°C. After three washes with PBS, samples were incubated with fluorescently labelled secondary antibodies (Alexa Fluor Plus, Thermo Fisher Scientific, or FITC #F-1005, AvesLab), diluted 1:1000 in blocking buffer, for 2h at RT. After three PBS washes, nuclei were stained for 20 min at RT with 1μg/ml 4′,6-diamidino-2-phenylindole (DAPI) in PBS. Thermo Fisher Scientific, #D1306). Mowiol 4-88 (Sigma-Aldrich, #81381) was used as mounting medium and the slides dried overnight before the acquisition.

### Confocal microscopy and imaging analysis

Confocal microscopy was performed by using Stellaris 5 confocal laser scanning system (Leica Microsystems), using a dry 20x or immersion 63x objective. Scale bars are reported in the figure legends. Confocal acquisitions were processed using ImageJ Software or Imaris Software (Bitplane).

#### Microglia density analysis

Microglial density was measured on z-stacks acquired at 20x magnification, and calculated based on co-localization of IBA1 and DAPI signals. DAPI and IBA1 signals were thresholded using fixed settings within the same experiment. DAPI signal was multiplied by IBA1 signal per each focal plane of the z-stack acquisition, using the “image calculator” function. The resulting mask was max-projected and the identified microglia nuclei were counted with the “analyze particle” function.

#### Microglial morphometric analysis

3D reconstruction was performed using Imaris Software (Bitplane), with the built-in Surface module on confocal z-stacks acquired at 63x magnification. IBA1 volume was quantified by applying 3D surface rendering of confocal stacks, using identical settings (fix thresholds of intensity and voxel) within each experiment. Surface and volume parameters were extracted for each individual reconstructed cell.

### Statistical analysis

We present all data as mean ± standard error of the mean (SEM) or median in violin plots. We performed statistical analyses using Prism 8.0 (GraphPad Inc., CA, USA), SPSS Statistics 26.0 (IBM Corp., NY, United States), Matlab R2022b (Natick, MA, USA), and R (version 4.3). Two-way repeated measures analysis of variance (RM-ANOVA, adjusted using the Greenhouse-Geisser correction when necessary) or mixed models were used to assess treatment effects on NREMS duration or fragmentation and SWA time courses. Significant hypothesis-driven main effects or interactions were subjected to post hoc assessments using Dunnett’s multiple comparisons tests. We analysed the behavioural performance by comparing each group’s average recognition index to chance level, a recognition index of 0.5, with a one-sample *t*-test. A significance level of *p* <.05 was used for all statistical analyses. For the analysis of histopathological measures, we used Bayesian approaches. The number of APP+ axonal bulbs was analysed using a Bayesian Negative Binomial Regression (brms package for R, version 2.19). The prior for this model was established with the coefficients resulting from a Poisson model as the expected values and a student distribution with the overall data’s standard deviation set as the dispersion value; this provided wide distributions and therefore a more naïve prior. To analyse MBP optical density values, we set up a Bayesian linear regression using a prior with coefficients from the expected values, obtained from a linear model, following a student distribution set as before. Spearman’s correlations were tested between SWA and number of APP+ axonal bulbs, and between SWA and MBP optical density values, as these relationships may not necessarily move in the same direction at a constant rate.

## Results

### SWA enhancement associated with improved memory upon upCLAS in TBI rats

Rats were subjected to either mock CLAS (non-TBI, TBI mockCLAS) or up-phase targeted CLAS (TBI upCLAS) for 5 days acutely after induction of TBI or sham (non-TBI) procedures, followed by 2 days of carry-over (no stimulation) EEG/EMG recording days (**Figure 1a**). Diagnostics data show that the vast majority of upCLAS-sound delivery successfully targeted the up-phase (0-90° targets) of ongoing slow waves during NREM sleep (**Figure 1b**), validating the accuracy of the methodology. Our results demonstrate that applying upCLAS to rats for 5 days immediately after TBI enhances overall SWA by in average 20.7%±4.9% in stimulation days 1-5 in respect to baseline (**Figure 1c**) without affecting the duration (**Figure 1d**) or fragmentation (**Figure 1e**) of NREM sleep. Moreover, we do not observe SWA rebounds upon discontinuing upCLAS.

Fourteen days after trauma (*t = 14*), i.e. nine days after CLAS paradigm discontinuation, we assessed whether acute delivery of upCLAS immediately after trauma alleviated sub-chronic posttraumatic cognitive impairment by comparing the performance of non-TBI rats and mockCLAS-treated TBI rats with that of upCLAS-treated TBI rats on the novel object recognition test (**Figure 2a**)^34^. As expected, non-TBI rats perform well at discriminating the novel object above chance level, whereas mockCLAS-treated TBI animals do not, reflecting impaired episodic memory. Rats treated with upCLAS immediately after TBI exhibit cognitive ability broadly equivalent to that of non-TBI rats (**Figure 2b**). The treatment with upCLAS sustains the beneficial effect previously shown to be conferred by pharmacologically enhanced SWA after TBI^21^, but avoids the potential off-target effects, altered consciousness, and unwanted tolerability and dependency concerns that are associated with drug-induced deep sleep.

**Figure 2.**
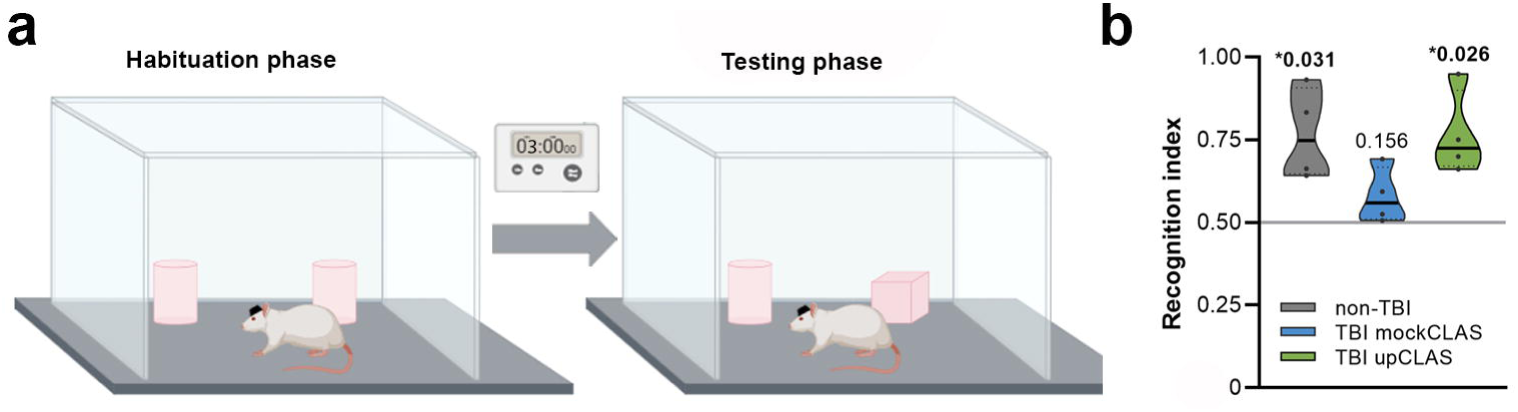
UpCLAS prevents cognitive decline in TBI animals. **a)** Novel object recognition test was used to probe non-TBI or mock/ upCLAS-treated TBI rats’ cognitive performance 14 days after trauma induction. **b)** Non-TBI rats perform significantly above chance level (n = 4, non-TBI, \**p* = 0.031) in the novel object recognition test, in contrast to mockCLAS-treated TBI rats (n = 4, TBI mockCLAS, *p* = 0.156), which showed a pronounced cognitive deficit 14 days post-TBI. TBI rats that received the upCLAS regime for 5 days acutely after trauma present above-chance-level recognition indices (n = 4, TBI upCLAS, \**p* = 0.026) 14 days after TBI. All groups were analysed by one-sample *t*-test in comparison with 0.50, defined as the chance performance value equivalent to ‘no recognition’. upCLAS: up-phase targeted closed-loop auditory stimulation; mockCLAS: flagging of triggers’ targets without sound delivery; TBI: traumatic brain injury, non-TBI: sham operated rats.

### Rescued histopathological damage upon upCLAS in TBI rats

In DAI, APP-filled bulbs that form in damaged axons can disrupt critical cortical-subcortical pathways, leading to widespread cognitive dysfunction and other neurological sequelae in both TBI patients and animal models^35,36^. To determine the extent of posttraumatic DAI upon CLAS-modulated SWA, we performed APP-specific immunostainings (**Figure 3a**) followed by stereological quantification of APP+ axonal bulbs in the anterior portion of the corpus callosum (**Figure 3b**), a particularly vulnerable white tract bundle, on coronal slices from brains harvested twenty-eight days after trauma (*t = 28*), i.e. twenty-three days after CLAS paradigm discontinuation. Comparison of Bayesian posteriors distribution (**Figure 3c**) showed substantially higher APP+ axonal bulb estimates for white matter tracts in the brains of mockCLAS-treated TBI rats than in non-TBI and upCLAS-treated TBI rats’ brains. This comparison demonstrates for the first time that nonpharmacologically enhanced SWA shortly after trauma leads to reduced posttraumatic secondary brain injury. Moreover, DAI levels, determined via number of APP+ axonal bulbs, positively correlate with stimulation-mediated changes in SWA measures (**Figure 3d**).

**Figure 3.**
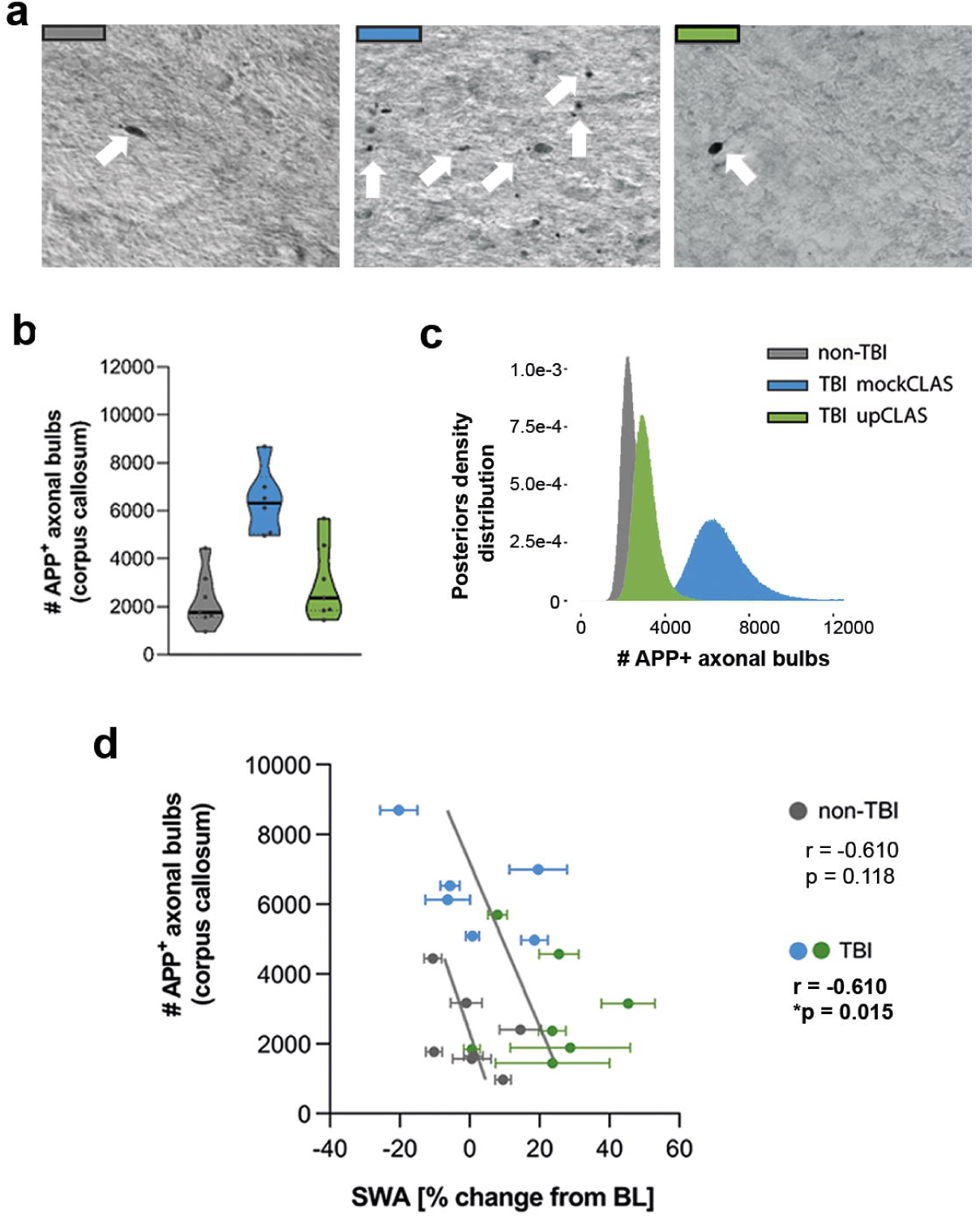
Reduced diffuse axonal damage in corpus callosum in TBI rats treated with upCLAS. **a)** Micrographs at 63**×** magnification display low or high burden of APP+ axonal bulbs (white arrows) in the corpus callosum of non-TBI (n = 7, grey), TBI mockCLAS (n = 6, blue), and TBI upCLAS (n = 7, green) rats coronal brain sections. **b)** Stereological estimates of the number APP+ axonal bulbs in corpus callosum of non-TBI, TBI mockCLAS and TBI upCLAS rats were plotted into violins and analysed by fitting them to a negative binomial distribution with treatment as categorical factor using a Bayesian general linear regression approach. The prior for this model was established with the coefficients of a Poisson distribution and following a student distribution. **c)** Histogram of the posterior density distributions for the parameters of each treatment. Strong differentiation of the TBI mockCLAS group, whose intervals stand clearly outside the 95%CI of both other groups. The TBI upCLAS and non-TBI posterior distributions have large superimposition with 13.04% of the posterior differences > 0 (non-TBI minus TBI upCLAS). **d)** Spearman’s correlation indicates a negative relationship between SWA (% of change from BL) and number of APP+ bulbs in the corpus callosum of TBI animals (*r* = 0.610, *\*p* = 0.015), whereas no correlation between these variables is observed in the non-TBI group. upCLAS: up-phase targeted closed-loop auditory stimulation; mockCLAS: flagging of triggers’ targets without sound delivery; CI: confidence interval; OD: optical density; BL: baseline; APP: amyloid precursor protein; SWA: slow-wave activity; TBI: traumatic brain injury, non-TBI: sham operated rats.

Furthermore, DAI-triggered demyelination represents one of the critical TBI-triggered secondary pathways to axotomy, eventual cell death, and consequent functional impairment^37^. Thus, we also assessed the extent of posttraumatic demyelination with specific immunostaining of the myelin basic protein (MBP) (**Figure 4a**), involved in myelin sheath formation and used in clinical settings to evaluate the severity of TBI^38,39^. We found that the corpus callosum of mockCLAS-treated TBI rats exhibited lower MBP staining levels than non-TBI controls as assessed by optical density quantification (**Figure 4b**), highlighting the damaging effect of TBI on principal fibre bundles in the rat brain. In contrast, the corpus callosum of upCLAS-treated TBI rats shows close-to-healthy MBP staining levels, with optical density values significantly overlapping (95% CI) with the non-TBI group (**Figure 4c**). Lastly, demyelination scores, determined via MBP optical density values, positively correlate with stimulation-mediated changes in SWA measures (**Figure 4d**).

**Figure 4.**
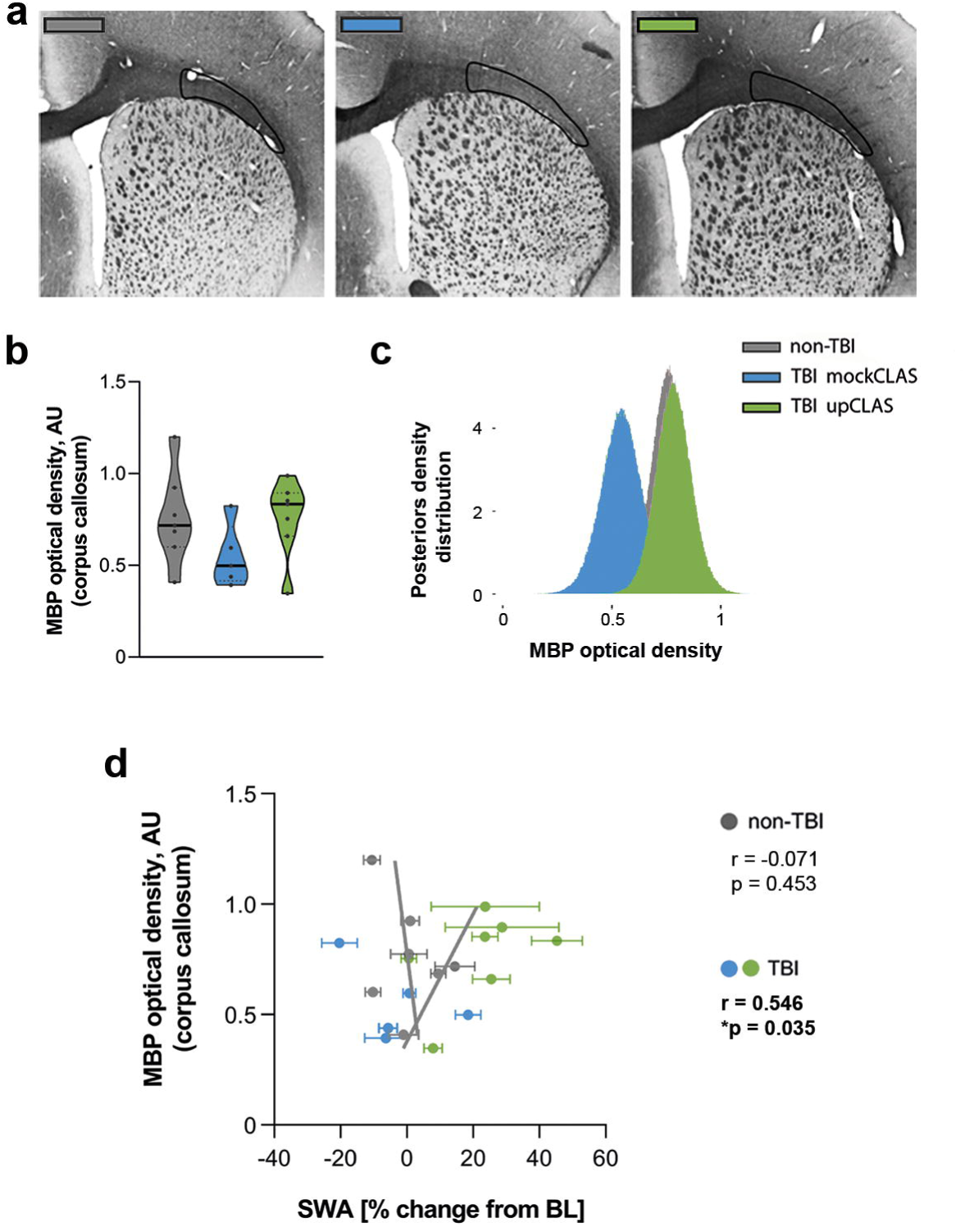
Reduced corpus callosum demyelination in TBI rats treated with upCLAS. **a)** Micrographs at 10**×** magnification displaying representative MBP staining levels in the corpus callosum of non-TBI (n = 7, grey), TBI mockCLAS (n = 5, blue), and TBI upCLAS (n = 7, green) rats coronal brain sections.**b)** OD values (AU) in the corpus callosum of each group were plotted into violins and analysed by a Bayesian linear regression fitted to robust priors (student_t in brms) for each group set according to their approximate normal parameters. **c)** Histogram of the posteriors’ density distribution for the parameters of each treatment. Comparisons showed an estimated difference in OD between TBI mockCLAS and TBI upCLAS of 0.24 ± 0.127 (AU), with 97.1% of the posterior differences > 0, compared to 60.4% when between non-TBI and TBI upCLAS groups. **d)** Spearman’s correlation indicates a positive relationship between SWA (% of change from BL) and MBP OD (AU) in the corpus callosum of TBI animals (*r* = 0.546, \**p* = 0.035), whereas no correlation between these variables is observed in the non-TBI group. upCLAS: up-phase targeted closed-loop auditory stimulation; mockCLAS: flagging of triggers’ targets without sound delivery; CI: confidence interval; OD: optical density; AU: arbitrary units, BL: baseline; MBP: myelin basic protein; SWA: slow-wave activity; TBI: traumatic brain injury, non-TBI: sham operated rats.

Combined, these histopathological results suggest a direct link between enhanced deep sleep and alleviated histopathological posttraumatic sequelae that presumably leads to decreased cognitive impairment.

### Enhanced microglial response upon upCLAS in TBI rats

Acute microglial reactivity after TBI may benefit the brain healing process by facilitating waste removal and clearance of cellular debris. Concomitantly, glia function has been linked to the sleep/wake cycle^15,17^. To investigate whether CLAS-mediated SWA enhancement in the posttraumatic acute phase triggered a differential microglial response in rat TBI brains potentially underlying the observed long-term ameliorated APP and MBP profiles, we explored IBA1 (microglia marker) fluorescent staining distribution in coronal non-TBI, TBI mockCLAS and TBI upCLAS brain sections 28 days after trauma (**Figure 5**). We found a visually distinct IBA1 distribution profile between the groups, with non-TBI brain presenting a homogenous scattering typical of homeostatic microglia (**Figure 5a**), as opposed to that observed in TBI upCLAS brains, where IBA1 fluorescence appeared clustered in localized areas of the brain, particularly noticeable in cortical structures (**Figure 5c**). TBI mockCLAS brains, on the other hand, presented an intermediate distribution profile (**Figure 5b**). We corroborated the visual observations via quantifications of microglia density (**Figure 5d**) and area covered by IBA1 (**Figure 5e**), demonstrating statistically significant differences between non-TBI and TBI upCLAS brains (One-way ANOVA, density: **p=0.0055; area covered: *p=0.044), but only observing non-significant trends between non-TBI and TBI mockCLAS groups.

**Figure 5.**
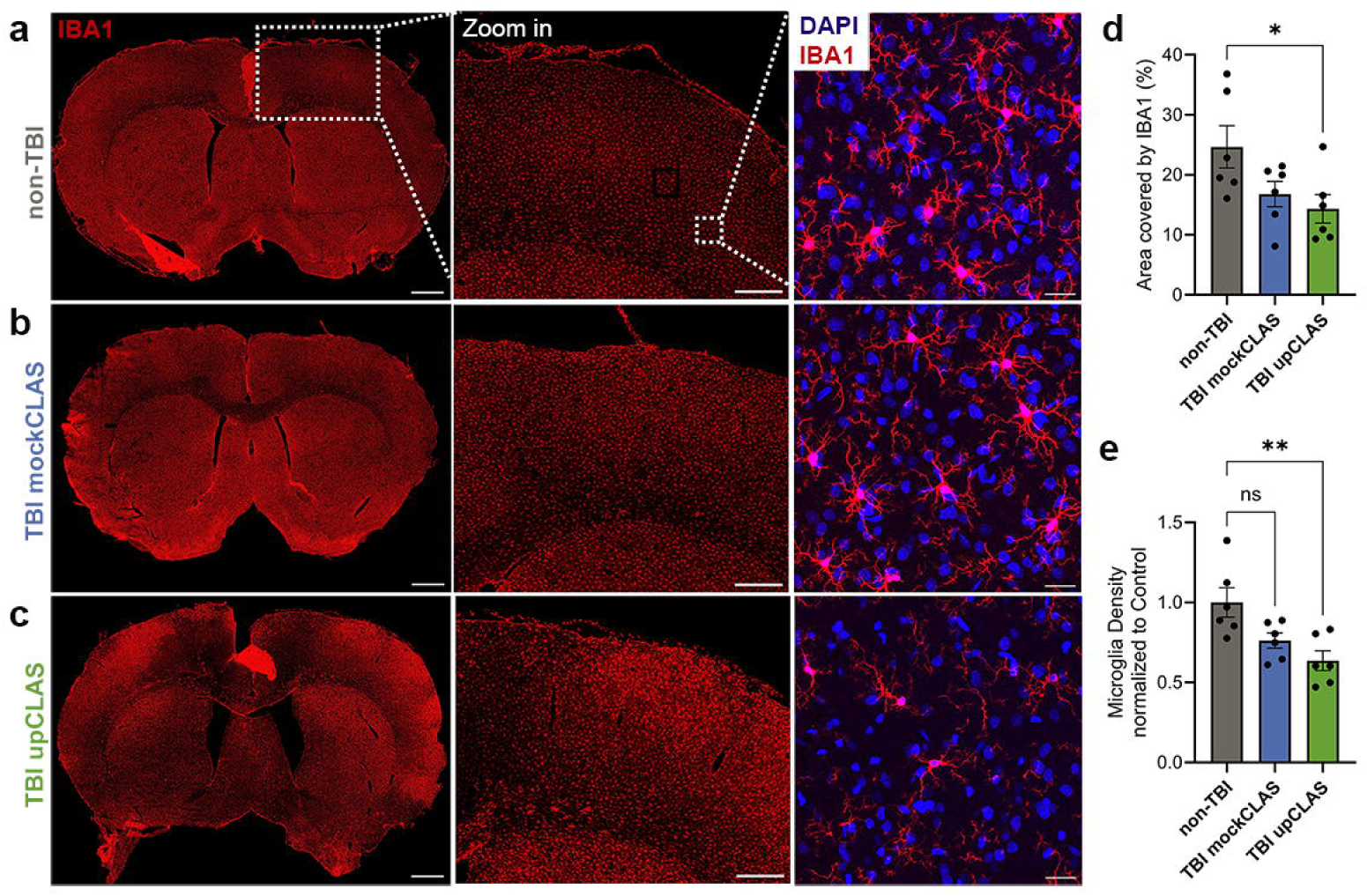
Clustered microglia distribution in upCLAS-treated TBI brains. **a)** Representative photomicrographs of IBA1 immunosfluorescence of 40µm coronal brain sections from non-TBI rats (n = 6), **b)** TBI rats treated with mockCLAS (n = 6), and **c)** TBI rats treated with upCLAS (n = 6) for 5 days upon injury. Distinct distributions are observed across the groups, as also observed in the zoomed in panels. High magnification shows IBA1^+^ microglia (red) and DAPI staining cell nuclei (blue). **d)** Quantification of cortical areas covered by IBA1^+^ cells in all three groups, shows reduced coverage in the TBI upCLAS animals (One-way ANOVA, non-TBI v. TBI upCLAS *\*p* < 0.05), evidencing a non-physiological distribution of microglia (reactive profile). **e)** Microglia density analysis also revealed a significantly lower number of IBA1^+^ cells/µm^3^ in upCLAS-treated TBI rat brains, characteristic of a patched distribution pattern. Bars represent 20 µm. upCLAS: up-phase targeted closed-loop auditory stimulation; mockCLAS: flagging of triggers’ targets without sound delivery; TBI: traumatic brain injury; non-TBI: sham operated rats.

Moreover, visual inspection of high magnification images (**Figure 5a-c**, rightmost panels, blue: DAPI; red: IBA1) from somatosensory cortical regions indicated that microglia morphology in the TBI upCLAS brains appear divergent to that of the other groups, with less processes and smaller size. To confirm these qualitative observations, we additionally performed microglia morphological analyses through 3D cellular reconstruction from high magnification micrographs of between 109 and 122 IBA1^+^ cells from n=6 subjects per group (**Figure 6a-c**), and compared derived quantitative measures of cellular area (**Figure 6d**) and volume (**Figure 6e**). We did not find significant differences in microglia area or volume between non-TBI and TBI mockCLAS brains, whereas microglia morphology in TBI upCLAS brain tissue was indeed significantly different from that of non-TBI animals (One-way ANOVA, area: **p=0.0046; volume: **p=0.0098), with smaller area and volume values suggesting a reactive profile.

**Figure 6.**
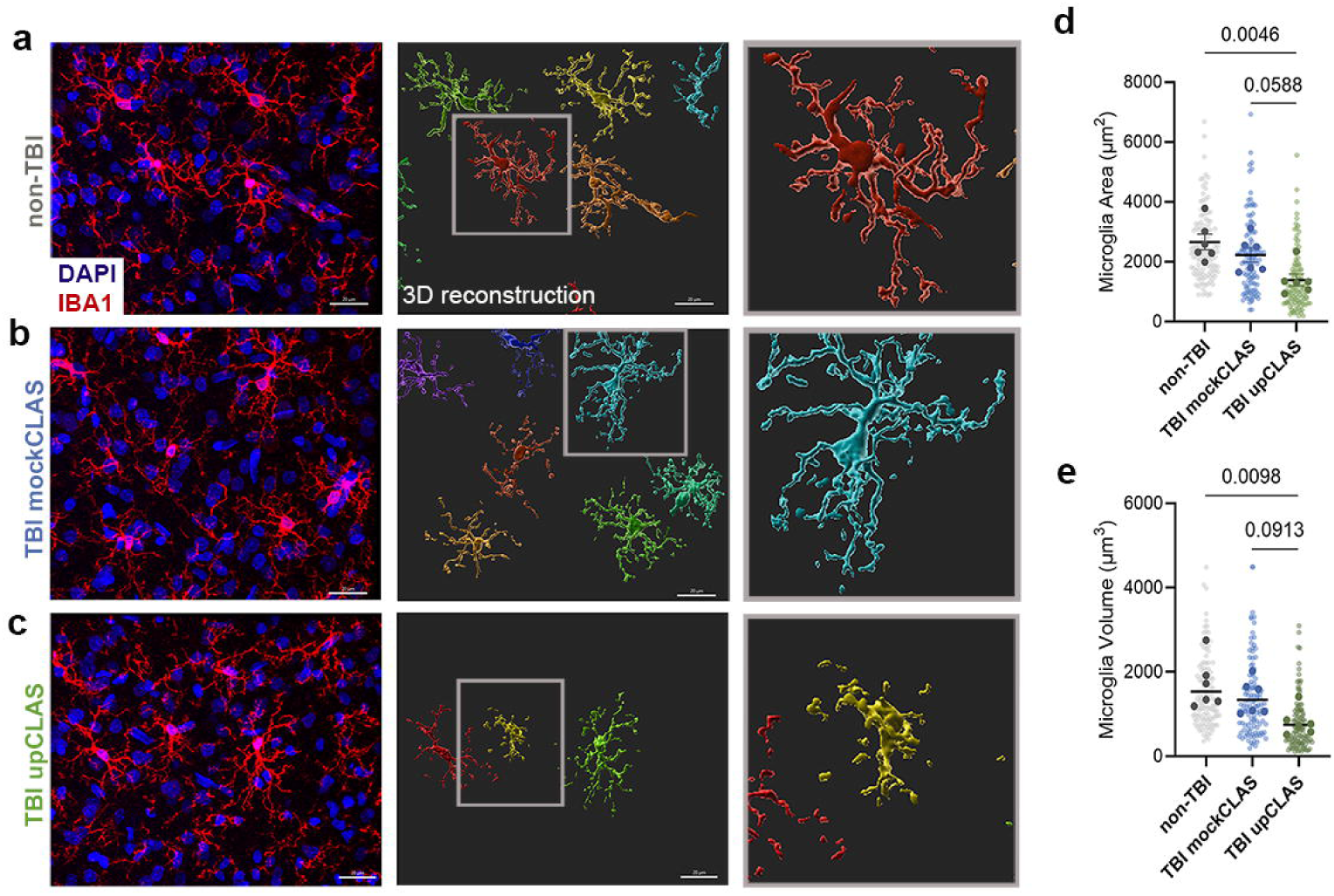
Responsive microglia morphology in upCLAS-treated TBI brains. **a)** Representative photomicrographs (left), Imaris-3D profile reconstructions (middle) and detailed representative 3D insets (right) of immunofluorescence-stained IBA1^+^ cells in somatosensory cortical regions of non-TBI (n = 122 total cells), **b)** TBI mockCLAS (n = 117 total cells), and, **c)** TBI upCLAS (n = 109 total cells) brains demonstrate differences between upCLAS-treated group and the others. **d)** 3D-reconstruction derived quantification of cellular area and **e)** volume in all three groups confirm qualitative observations, demonstrating that TBI upCLAS animals present significantly lower values in both measures than non-TBI rats (One-way ANOVA, Tukey’s multiple comparisons test: non-TBI v. TBI upCLAS, area: *\*\*p* = 0.0046; volume: *\*\*p* = 0.0098), indicating a more responsive microglia profile in SWA-enhanced rat brains. Trends were observed towards lower area and volume values in TBI upCLAS group compared to TBI mockCLAS (One-way ANOVA, Tukey’s multiple comparisons test: TBI mockCLAS v. TBI upCLAS, area: *p* = 0.0588; volume: *p =* 0.0913), suggesting TBI mockCLAS brain to present intermediate values between non-TBI and TBI upCLAS groups. Bars represent 20 µm. upCLAS: up-phase targeted closed-loop auditory stimulation; mockCLAS: flagging of triggers’ targets without sound delivery; TBI: traumatic brain injury; non-TBI: sham operated rats; 3D: tridimensional.

Together, these results indicate microglial response was boosted by upCLAS treatment in TBI rats in the acute phase, thus evidencing a still non-resumed reactivity profile in the chronic period 28 days after trauma.

## Discussion

The present results demonstrate that upCLAS effectively enhances SWA, reduces DAI, preserves myelin integrity, and mitigates posttraumatic memory impairment in TBI rats, along with an increased microglia reactivity. To our knowledge, this is the first report demonstrating an overall beneficial effect of a nonpharmacological and noninvasive sleep-based treatment approach on structural brain integrity following TBI, together with first insights into a potential underlying mechanism.

Multiple studies in humans and animals have investigated the effects of CLAS on boosting^40–44^ and disrupting^45,46^ SWA. So far, however, no report has evidenced the effectiveness of CLAS in the context of brain damage. Here we demonstrate the efficacy of CLAS in rodent TBI by showing upCLAS is feasible, safe, stable, specific, and effective in enhancing SWA in TBI rats over multiple days, while not altering deep sleep duration or fragmentation. Analysis of carry-over effects on EEG spectral power in NREM sleep revealed a renormalization of SWA upon CLAS’ discontinuation. Of note, however, renormalized levels of SWA in upCLAS TBI rats do not resume to non-TBI levels (see Figure 1a). Moreover, SWA levels in mockCLAS TBI rats appear to go naturally up starting day 4-5 of the protocol. This is consistent with previous report of transiently increased SWA ∼1 week after trauma induction in this TBI model^47^. Therefore, the confluence of the mockCLAS TBI and upCLAS TBI SWA curves during the carry-over days implies a complete renormalization of SWA values back to their ‘new’ baseline level upon upCLAS discontinuation. This data is consistent with findings reported in human CLAS^48^ and denotes a desirable high degree of reversibility. These findings open valuable new avenues for translation into clinical therapy avoiding pharmacological SWA enhancement after TBI, which is hindered by ethical and practical limitations. In addition to their dependency, tolerance, and sleep-architecture-altering issues, sleep-inducing pharmacotherapies impede swift clinical monitoring of patients after TBI. From an EEG standpoint, a limitation of our work consists of that we implanted only two diametrically opposed EEG channels per rat, equidistant from the TBI site, which does not suffice for analysis of changes in functional connectivity and network geometries in TBI brains’ response to CLAS. Future experiments using LFP/multiunit activity multi-electrodes could provide a more comprehensive understanding of CLAS’s influence on surface and deep brain network remapping and posttraumatic global and local structural connectivity.

Both intra- and extracellular mechanisms may be responsible for the reduction in APP accumulation in axonal bulbs following upCLAS-enhanced SWA. A widely discussed possibility is that enhanced glymphatic function facilitates removal of proteinaceous products during sleep^49,50^, and upregulation of protein players within other processes has been proposed, such as the ubiquitin/proteasome system^51^. However, direct links remain to be fully established between these homeo-/proteo-static pathways and TBI-triggered pathomechanisms, with focus on whether such systems are truly able to facilitate clearance of high molecular weight proteins and cellular debris. Moreover, TBI-induced secondary injury extends beyond abnormal protein accumulation due to impaired axonal transport^52^. For instance, TBI-triggered proteinolysis mediated by calpain activation after axonal injury often leads to demyelination^53^. Our finding of preserved MBP staining intensity in the corpus callosum - the main white matter tract in the rodent brain - indicates conserved myelination levels in upCLAS-treated TBI rats and thus suggests that acute auditory enhancement of sleep SWA may be necessary and sufficient to promote neuroprotection after TBI.

Improved posttraumatic cognitive ability upon auditory slow-wave sleep enhancement nine days after stimulation discontinuation is a compelling finding. However, these results must be interpreted with caution. The high variability of NORT scores across all experimental groups required us to perform a median-split analysis to identify and exclude a subset of rats in each group that presented very low or no behavioural engagement in the test (data for non-engagers, similar for all groups, not shown)^34^. Considering that a similar number of rats was excluded from each group, we assume the inactivity of these subjects was not related to the treatment and surmise that these non-engagers were particularly susceptible to poor experimental conditions, such as environmental noise and odours. Nonetheless, our preliminary analysis of behavioural scores from rats that actively engaged in the test (engagers, Fig. 2) strongly suggests that acute enhancement of SWA after TBI helps preserve cognitive abilities in the mid-term.

In association with alleviated TBI outcomes at functional (*t = 14*) and neuropathological levels (*t = 28*) in the sub-chronic phase post-trauma, we also observed an altered microglial profile in upCLAS-treated brains, where more reactive IBA1 distribution profile and morphology were observed. These key results suggest that enhanced SWA likely heightened microglia reactivity in the acute phase after trauma LJ this not having resumed back to baseline levels at *t = 28* LJ that positively conditioned lessened sequelae. The exact mechanisms at play remain to be determined. It has been recently shown that microglia Ca^2+^ activity is naturally higher during sleep than wakefulness^54^. Growing evidence link Ca2+ activity in microglia to enhanced phagocytic capacity^55,56^. Thus, it is conceivable that acute post-TBI treatment with upCLAS promotes phagocytic activity in the injury site, known to contribute to clearance of debris and release of neurotrophic factors that support neuronal survival and recovery^57^. However, further in-depth experiments are necessary to fully understand the role of SWA enhancement onto microglia response profiles and their temporal timing, including determination of cytokines levels to concretely characterize the brain’s inflammatory state.

On the other hand, microglia have been recently postulated to be involved in hippocampal synaptic transmission in relation to regulation of sleep duration during the light/dark cycle. In fact, microglia depletion can disrupt sleep/wake cycles and circadian rhythms^58^, suggesting a tight interplay between this crucial cellular population of the neuro-immune response and sleep quality/ quantity. Alterations of the circadian rhythms in microglia can deeply affect the immune response, phagocytic function, metabolism, and other aspects of microglia, which play a key role in neurological diseases^17,59^. In this line, implementation of upCLAS increasing the intensity of deep sleep (SWA) in a mouse model of Alzheimer’s disease, partially rescued the intrinsic circadian misalignment observed in the light-dark transition in these mice^60^, suggesting an indirect role of upCLAS-mediated SWA enhancement on circadian rhythm strengthening. Therefore, it is plausible that CLAS-mediated SWA enhancement in TBI rats facilitated a more efficient sleep pressure dissipation thus reinforcing their circadian/sleep-wake cycle, with consequent up-regulation of the microglial response.

Conversely, mockCLAS-treated TBI animals exhibited microglial morphology and density akin to non-TBI animals at *t = 28*. This microglial phenotype may be attributed to a resolved acute inflammatory response after TBI followed by subsequent return of this cell population to a more homeostatic state. In fact, microglial response peaks shortly after injury, followed by a gradual decline in reactivity markers^61,62^. In the absence of acutely sustained SWA enhancement, the microglial population in the mockCLAS-treated TBI brains may have reverted at *t = 28* to a less responsive state, characterized by a ramified morphology and baseline-like density around the injury site. Overall, our observations suggest that acute microglia reactivity may have enabled mitigating posttraumatic DAI and demyelination in the sub-chronic posttraumatic phase in upCLAS-treated TBI animals which might reflect a role of SWA to promote a pro-regenerative state of microglial cells, potentially enhancing their clearance activity, fostering survival and repair mechanisms.

Follow-up work shall carefully explore whether the presently observed positive effect on microglia response upon upCLAS remains true in chronic TBI stages (3, 6 or 12 months) and/or in the presence of secondary stressors.

Altogether, the apparent ability of upCLAS-mediated SWA enhancement to influence the trajectory of microglial response suggests that early deep sleep interventions may be key in altering histopathological and behavioural recovery after TBI. In this scenario, auditory stimulation seems to facilitate a more favourable microglial microenvironment aiding positive resolution of histopathological sequelae and consequent mitigation of long-term TBI symptoms^63^, in line with our results on spared cognitive performance in upCLAS-treated TBI rats. Elucidating the exact mechanisms by which deep sleep enhancement regulates restoratives processes involving, perhaps not only, modulation of the type and temporal kinetics of microglial reactivity, with particular focus placed on the acute (1-7 days) and chronic phases (3-12 months) of the neuro-immune response, shall be a main objective of follow-up studies aimed at determining the safety and efficacy of CLAS implementation as therapeutic strategy in TBI. In addition, these findings offer an attractive platform for testing the effect of nonobtrusive and nonpharmacological deep sleep enhancement on clinical and radiological outcomes in TBI patients.

In summary, acute posttraumatic delivery of upCLAS is the first nonpharmacological, noninvasive therapeutic sleep-based tool to significantly prevent TBI-related histopathological sequelae and thus alleviate cognitive deficits, presumably via beneficially boosted microglial response. Our data strongly encourages not only further preclinical investigations into the mechanisms governing the associations we observed, but meticulous translation of this technology into clinical environments to test its safety and efficacy in TBI patients.

## Funding

This project was funded by the Swiss National Science Foundation (grant nr. 163056, CRB), the Clinical Research Priority Program ‘Sleep and Health’ of the University of Zurich (CRB), the SleepLoop Flagship project of Hochschulmedizin Zürich (CRB), the University of Zurich (Einrichtungskredit, CRB), the Neuroscience Center Zurich by donation of Rahn & Bodmer banquiers (DN), the Dementia Research – Synapsis Foundation Switzerland with an earmarked donation from the Armin & Jeannine Kurz Stiftung (DN); the ERC StGrant REMIND 804949 and funding from the University of Lausanne (RCP).

## Acknowledgments

We thank Mr. Mark Hanus, Mr. Myles Billard, Dr. Virginia Meskenaite, Dr. Sedef Kollarik, and Dr. Joachim Buhmann’s group for their time and valuable technical support.

## Notes

### Competing Interest Statement

The authors have declared no competing interest.

